# Using null models to infer microbial co-occurrence networks

**DOI:** 10.1101/070789

**Authors:** Nora Connor, Albert Barberán, Aaron Clauset

**Affiliations:** Department of Computer Science, University of Colorado, Boulder CO, USA.; Cooperative Institute for Research in Environmental Sciences, University of Colorado, Boulder, CO, USA.; BioFrontiers Institute, University of Colorado, Boulder, CO, USA.; Santa Fe Institute, Santa Fe NM, USA.

## Abstract

Although microbial communities are ubiquitous in nature, relatively little is known about the structural and functional roles of their constituent organisms’ underlying interactions. A common approach to study such questions begins with extracting a network of statistically significant pairwise co-occurrences from a matrix of observed operational taxonomic unit (OTU) abundances across sites. The structure of this network is assumed to encode information about ecological interactions and processes, resistance to perturbation, and the identity of keystone species. However, common methods for identifying these pairwise interactions can contaminate the network with spurious patterns that obscure true ecological signals. Here, we describe this problem in detail and develop a solution that incorporates null models to distinguish ecological signals from statistical noise. We apply these methods to the initial OTU abundance matrix and to the extracted network. We demonstrate this approach by applying it to a large soil microbiome data set and show that many previously reported patterns for these data are statistical artifacts. In contrast, we find the frequency of three-way interactions among microbial OTUs to be highly statistically significant. These results demonstrate the importance of using appropriate null models when studying observational microbiome data, and suggest that extracting and characterizing three-way interactions among OTUs is a promising direction for unraveling the structure and function of microbial ecosystems.

**Author Summary:** Microbes are ubiquitous in the environment. We know that microbial communities – the groups of microbes that live together, interact, and depend on one another – vary across environments. Multiple processes, ranging from competition between microbes to environmental stress, are believed to alter microbial community composition. Here, we describe a set of statistical techniques that can more accurately identify the underlying taxa relationships that structure the observed abundances of microbes across habitats. Using a large data set of soil samples collected across North and South America, we both illustrate the statistical artifacts that incorrect methods can introduce and describe proper techniques based on appropriate null models for studying how the abundances of taxa vary across soil samples. These tools improve our ability to distinguish ecologically meaningful interactions from simple statistical noise in such observational data. Our application of these tools suggests some previous claims about the network structure of microbial communities may be statistical artifacts. Furthermore, we find that three-way interactions among microbial taxa are significantly more common than we would expect at random, and thus may provide a novel means for identifying ecologically meaningful interactions.

## Introduction

Microbes play essential roles in many, if not most, ecosystems. They play particularly important roles in regulating agricultural systems (e.g. Navarrete *et al.* 2015), human health (for a review, see Cho & Blaser 2012), and may even have an effect on mental health and behavior (Yano *et al.* 2015). Yet despite the importance of microbes and the recent technological advances in the field, essential questions remain about the composition and ecological structure of these microbial communities. For instance, how do communities change in response to internal dynamics and external perturbations, and how could we design communities with novel functionality? Deeper insights into the variables that shape the structure and function of microbial communities would have wide-ranging significance, both practical and theoretical.

One difficulty in scientifically addressing questions about microbial communities comes from the inability to culture the vast majority of microbes in a laboratory environment (Rappe & Giovannoni 2003). Instead, microbial community composition must be inferred from sequence data obtained by environmental DNA sampling. This limitation restricts our ability to test for causal mechanisms that drive a microbial community’s structure and composition. Instead, observational data is often drawn from multiple samples across time or habitats (Barberán *et al.* 2015, Faust & Raes 2012, Peura *et al.* 2015, Steele *et al.* 2011, Kara *et al.* 2013). Complicating these efforts is a lack of robust statistical methods for analyzing these observational data in a way that reliably controls for plausible sources of variability and the spurious co-occurrence network patterns they can produce. Here, we present and test methods for extracting statistically significant co-occurrence patterns among microbes and for interpreting the induced network structure.

A common design for a microbial community observational study has the following form. Using high-throughput sequencing technologies, genetic data is extracted from a set of locations, such as soil, water, or host-associated habitats including fecal samples or cheek swabs. The observed DNA sequences are then binned into operational taxonomic units (OTUs), which are taxonomic categories for microbes and are based on a DNA sequence similarity threshold (usually 97% for 16S rRNA gene). This step is necessary due to the difficulty in objectively defining microbial species, since these taxa reproduce asexually and many have the ability to transfer genes horizontally. The OTUs are placed into an abundance matrix ***A***, where each element ***A***_*i,j*_ gives the number of sequences representing a particular OTU *i* observed in a particular sample or location *j.* This matrix is then used to identify pairwise interactions, under the assumption that OTUs whose abundances correlate across samples are likely to be ecologically related, either symbiotically or through similar environmental preferences. To obtain correlation values, a similarity measure is computed for each pair of vectors of OTU abundances across locations (Faust & Raes 2012), and statistically significant similarities are interpreted as potential ecological interactions. The set of such pairwise interactions among the sampled OTUs can be transformed into a network of microbial interactions, where nodes are OTUs and significant pairwise correlations are represented as edges in the network. This network’s structure can then be used to understand the community’s organization and function.

Such microbial interaction networks have many uses, not the least of which is making complex data visually interpretable. They also facilitate the investigation of underlying ecological processes that shape microbial communities. Past work on microbial networks has examined many of their structural properties, including an OTU’s degree (number of connections), an OTU’s betweenness centrality (a geometric measure of its network position), the network’s frequency of three-way interactions (the clustering coefficient), and the network’s average path length (a measure of system compactness). These properties have been measured for networks derived from a variety of habitats, including soil (Barberán *et al.* 2012), marine (Steele *et al.* 2011), and freshwater communities (Kara *et al.* 2013). For instance, nodes in a network that have high degree or high centrality may be interpreted as keystone taxa (Steele *et al.* 2011, Berry & Widder 2014, Williams *et al.* 2014). Recent work has shown that these keystone taxa play important roles in structuring microbial communities in plant-microbe interactions (Agler *et al.* 2016). A group of OTUs that tend to co-occur may correspond to taxa that share an ecological niche due to habitat filtering, or that participate in a symbiotic interaction (Faust & Raes 2012). Similarly, groups of OTUs that tend to mutually exclude each other may represent competitive interactions within a given niche. We may also compare the structure of these microbial communities with that of other biological networks (Williams *et al.* 2014), e.g., in order to understand whether principles from macroecology also hold for microbial communities.

Network structure can also shed light on how a microbial community may respond to environmental perturbations. A right-skewed degree distribution among OTUs may be evidence for robustness to high levels of random removal of species, or sensitivity to the targeted removal of the keystone taxa (Faust & Raes 2012, Peura *et al.* 2015). This network property may be related, for instance, to predicting whether a person’s gut microbiome will recover after a course of antibiotics. Similarly, network structure can facilitate the identification of community assembly processes, for instance, by comparing the structural signatures of neutral processes where all taxa are demographically equivalent, versus those produced by niche-structured processes like niche partitioning and competitive exclusion (O’Dwyer *et al.* 2012, Levy & Borenstein 2013, Pholchan *et al.* 2013, Tucker *et al.* 2015). Greater insight into assembly dynamics may facilitate predictions of community response to natural or artificial perturbations (Faust & Raes 2012).

The broad importance of microbial interaction networks makes it essential that they be reliably and accurately extracted from OTU abundance matrices, and that patterns in the resulting network structure be properly interpreted. However, within the standard approach to extracting these networks from co-abundance matrices are underlying statistical assumptions that can contaminate the network with spurious or misleading patterns. Specifically, spurious patterns in microbial co-occurrence networks may arise from matrix sparsity, the choice of correlation function, and the use of thresholds. Separate problems may arise when abundance data is normalized, making it compositional. Addressing the issues of compositional data is beyond the scope of this paper; however, in our conclusions we offer a brief discussion of their relationship to the methods described here. In the following sections we examine the consequences of spurious patterns in the data and leverage the ensuing errors as a motivation for the use of null models as the foundation for the statistical methods we introduce. Our methods are statistically principled methods, being based on standard null models, and allow us to more accurately distinguish ecological signals from statistical noise, both in the abundance matrix itself and in the distribution of edges in the derived network.

We demonstrate these techniques using a previously studied soil microbiome data set from North and South America (Barberán *et al.* 2012). We find that some measures of network structure are barely distinguishable from random noise, while others are more plausibly the result of ecological interactions. A notable example of the latter category is the network’s clustering coefficient, the density of three-way OTU interactions, which remains statistically significant when compared to each of our null models. We close with a brief discussion of the utility of null models in studying observational data and the ecological significance of triangles and modularity in microbial co-occurrence networks.

## Results

### Two classes of null models

Null models are a standard statistical approach for reliably identifying data patterns that cannot be attributed to simple sources of random variation. Data distributions that differ from a null model are thus potentially derived from complex processes. In our case, large deviations may be interpreted as potentially caused by ecological processes. One example of a null model is the common test of statistical significance, wherein we measure the likelihood of observing, under the null model, a particular statistical value or one more extreme. This probability is quantified by a standard *p*-value which has a uniform distribution when the true data generating process is the null model. Common choices for null models focus on a set of independent draws from a simple parametric distribution, e.g., flipping coins or rolling dice. Null models can be substantially more complicated, and in this case, numerical methods are typically required to calculate the null distribution of the test statistic. If a null model is chosen well, meaning that it incorporates plausible sources of random variation in the data, and the computed *p*-value still low (typically below the conventional but nevertheless arbitrary threshold of 0.05), then a deviation between the model and the data can indicate the presence of scientifically meaningful processes.

Here, we describe and study two classes of null models for inferring ecological interactions from a matrix of OTU abundances. The first class facilitates the extraction of significant pairwise interactions from the matrix in order to obtain a network. The second class facilitates the detection of significant patterns in the distribution of edges within the derived network.

In the rest of this section, we will introduce the first class of null models, in which we will incorporate existing variability in the observed data to identify pairwise interactions among OTUs. First, we correct the behavior of the Spearman rank correlation coefficient when the OTU matrix is sparse by breaking ties randomly. Second, in order to choose a threshold for significant interactions, we use matrix permutations to generate artificial matrices with the same naturally high variance as the data but which lack the correlations that are generated by ecological processes. Applying the tie-breaking step to these artificial matrices yields a null distribution of correlation scores, which provides a simple means for selecting a threshold for statistically significant interactions. If any pair of OTUs in the tie-breaking model has a correlation score above this threshold, we call this interaction statistically significant and include it in the interaction network; any correlation below the threshold is discarded.

In the second class of null models, we ask whether particular statistical patterns in the distribution of these interactions across the network are likely the result of random connectivity, and thus unlikely to be caused by ecological processes. Our approach here builds on standard random graph models from network science, which control for the average degree or the distribution of these degrees in order to construct an appropriate null distribution for other network properties. Characteristics that are independent of size and connectivity indicate co-existence of taxa, which may plausibly be attributed to ecological interactions or functions.

The fact that some properties can be explained by the size, degree, or connectivity of the network does not make them ecologically unimportant. In fact, the ecological impact of overall biodiversity as well as co-occurrence patterns (i.e., functional redundancy) is well established (Van Der Heijden *et al.* 2008, Philippot *et al.* 2013). In practice, these null models can be used to identify more complicated statistically interesting patterns, such as heterogeneous interactions among groups of microbes, that may relate to other ecological processes, either known or unknown.

### The abundance matrix of microbial soil communities

To illustrate the importance of examining microbial abundance data with respect to the two null model classes, we apply these methods to previously collected data on soil microbes sampled from 151 sites in North and South America (Lauber *et al.* 2009). From soil samples, Barberán *et al.* extracted 16S rRNA sequences and binned them into OTUs at a 90% rRNA sequence similarity threshold. They assigned taxonomy to OTUs using RDP Classifier (Wang *et al.* 2007) against the Greengenes database (DeSantis *et al.* 2006). To obtain the abundance matrix, they computed the number of sequences that mapped to each OTU at every sample site. To control for sample contamination and potential sequencing errors, they discarded OTUs with fewer than 5 sequences across all locations, which reduced the number of OTUs from 4,087 to 1,577.

Like many environmental DNA surveys, the resulting soil microbiome abundance matrix is very sparse. Abundance values of zero comprise fully 85% of the matrix. Most sites contained 150-300 OTUs, but only 1% of matrix entries have more than 10 sequences for a given OTU at a given site. In other words, although there were on the order of 1000 sequences from each location, most OTUs at a site were phylogenetically distinct.

In order to calculate the correlation of abundance patterns between a pair of OTUs, we must choose a similarity score function. The most common choices in past studies are Pearson and Spearman correlations, which exhibit good statistical sensitivity and specificity under standard conditions (Berry & Widder 2014). However, the Pearson correlation assumes that variables are normally distributed and linearly correlated, and it behaves poorly when relationships are nonlinear, as may be the case in complex microbial systems. Spearman’s rank correlation, which measures the degree to which two variables monotonically co-vary, does not suffer from this problem and is the more common choice in microbiome studies (Lozupone *et al.* 2012; see also Weiss *et al.* 2016 for a review of correlation methods).

### A correction for matrix sparsity in Spearman ranks

In this setting, Spearman will overestimate correlations when nearly all abundances are either zero or some integer close to zero. As an intermediate step, Spearman assigns a rank value to each location, and locations with equal abundance receive the same rank. Thus, both matrix sparsity and a heavy-tailed distribution of abundances will induce a very large number of multi-way ties, which will then have identical ranks. The result is an inflated pairwise correlation score under Spearman. (Standard implementations of Spearman’s in Matlab, R, and Python all rely on the user to correct for ties in the data.)

This behavior can be corrected through breaking ties at random by adding a small amount of real-valued noise to each entry in the abundance matrix. After adding these minor perturbations, the set of all pairwise Spearman rank correlation coefficients (*ρ*) form a smooth distribution (Figure 1A), as desired, rather than a perverse disjoint distribution when ties are not broken (Figure 1B).

**Fig 1:**
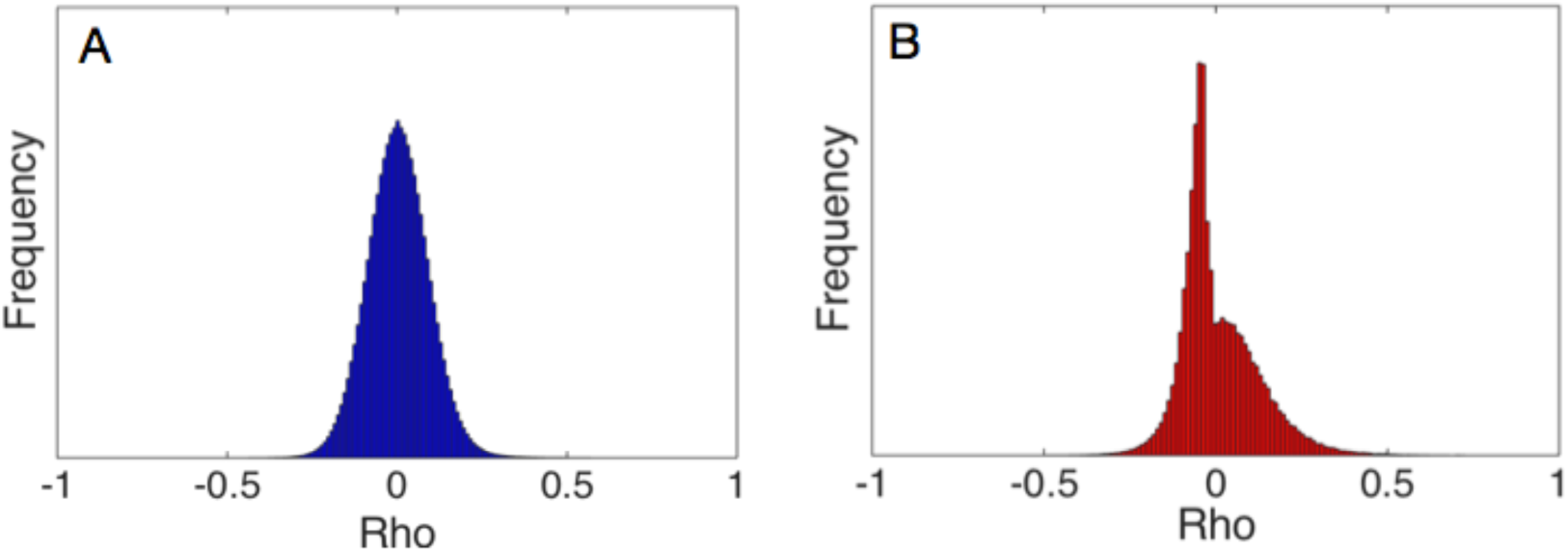
Null distributions of Spearman rank correlation coefficients across sites for the Barberan et al. soil microbiome data. (A) Coefficients under Monte Carlo sampling, using noise to break ties randomly. (B) Coefficients without correcting for tied ranks between locations.

Crucially, the noise added to each observed value must not disturb the partial ordering obtained without the noise. In practice, this is easily accomplished by using Monte Carlo to sample from the many total orderings that are consistent with the original partial ordering. Under a particular choice of significance threshold, this procedure will generate a set of equally plausible networks, which are free from the statistical artifacts of tied ranks.

This correction prevents the spurious conclusion that two taxa are ecologically related because they are both absent from many of the same locations. There are many reasons why a taxon could have zero abundance at a given location, including habitat filtering, local extinction due to ecological drift, dispersal limitation, or competitive exclusion. Or, it may indicate that the taxon’s DNA failed to bind to the 16S primer during amplification, was undetected due to sequencing depth, or was absent by chance from the soil sample. In short, an abundance of zero is highly ambiguous, and a conservative approach is to avoid inferring the presence of an interaction based primarily on shared absences.

### Converting the sparsity-corrected data into a network

To convert the abundance matrix into a network, we must apply a threshold to the similarity scores. In this way, only OTU pairs for which the absolute value of their score is above the threshold are connected in the network. It follows that a node with no scores above the threshold will have a degree of zero in the network, and by convention we omit such singletons from subsequent analysis (Barberán *et al.* 2012). As a result, the number of nodes *n* in the inferred network will typically be less than the number of OTUs *N* in the abundance matrix.

### Picking a threshold for significance

Choosing an appropriate threshold of significance for similarity scores is an open question, particularly for sparse data sets like OTU abundance matrices (Thomas & Blitzstein, 2011). The goal of this choice is to eliminate pairwise interactions that are likely due to statistical fluctuations or sampling noise, without excluding interactions due to biological processes. Furthermore, we would like the scientific conclusions that we draw from the resulting data to be robust to reasonable variations in threshold choice (Thomas & Blitzstein, 2011). Currently, however, there is no generally reliable method for balancing these two conflicting goals in OTU abundance matrices. Some studies have used random permutations of the abundance matrix to compute a null distribution of similarity scores, and then selected as a threshold the similarity value corresponding to a conventional *p*-value choice of 0.01 or 0.05 (Faust & Raes 2012). However, this procedure tends to select very low thresholds, and this may potentially result in a high false positive rate for interactions. Other studies have used arbitrarily chosen thresholds (Friedman & Alm 2012, Qin 2010).

Here, we use a repeated element-wise random permutation of the noise-added abundance matrix to first compute a null distribution of similarity scores. We then compute the size of the largest component -- the largest set of nodes for which any pair is connected by some sequence of edges -- in the induced network for a wide range of threshold values. Because the permutations break any ecologically-driven correlations in the abundance matrix, this curve has a characteristic sigmoidal shape (Figure 2). The location of the curve’s transition to less than 1% of OTUs in the largest component serves as a reasonable choice for the lower bound on the threshold. Networks derived from this permuted data treatment are composed of all spurious links, so a threshold below that transition, which would include these links, is overly inclusive. In practice, a conservative choice of threshold will be a value slightly above this transition point. Including the sparsity correction from above within this procedure serves to correct the substantial distributional bias in similarity scores that would otherwise occur (see Figure 1) as a result of multiple tied ranks and the heavy-tailed distribution of abundance values.

**Fig 2:**
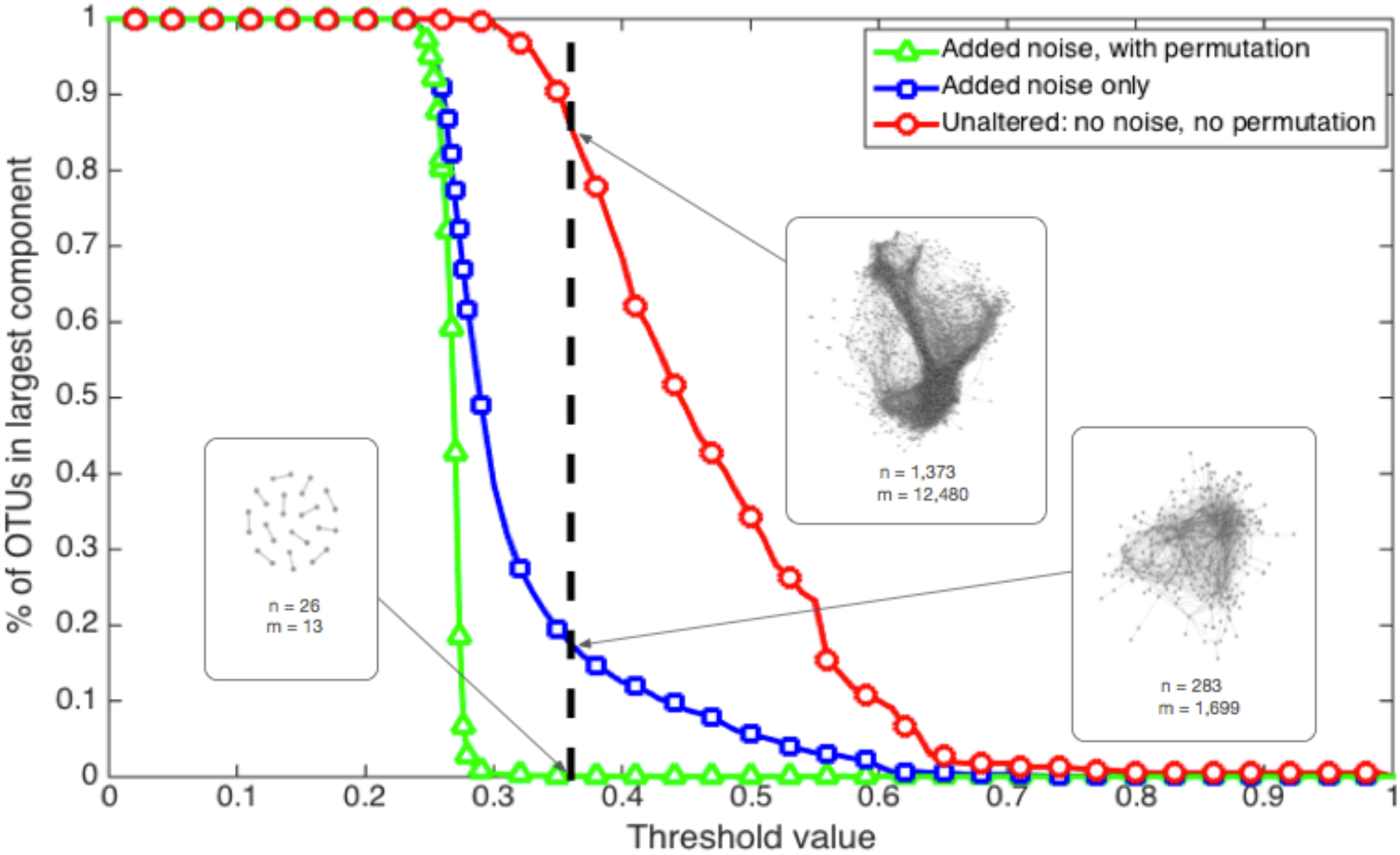
Fraction of all OTUs in the largest component, as a function of correlation threshold. When the pairwise correlation threshold is 0, all edges are included and thus all nodes are in the largest component. When the threshold is 1, all edges are excluded and all singletons are discarded, so all of the OTUs are excluded from the analysis. The inset networks result from applying a threshold of 0.36, shown by the bold dashed line, to each of the treatments. The 0.36 threshold corresponds to 86% of OTUs in the largest component for the unaltered data, but just 18% of the OTUs in the noise-added treatment. For the permuted treatment, with noise added, the threshold intersects after the phase transition, yielding <1% of OTUs in the largest component.

We subject the OTU abundance data to three different treatments and systematically vary the threshold to illustrate its impact on each. The three treatments are (i) the original data, (ii) the original data with the Spearman correction, and (iii) the original data with both Spearman correction and permutation null distribution. To illustrate the effect of threshold choice on each treatment, we measure the fraction of OTUs *N* contained in the largest component of the network across similarity thresholds (Figure 2). The size of this component provides a simple quantitative measure of overall graph connectivity, and is a monotonically decreasing function of the threshold. That is, higher thresholds will tend to produce smaller, less connected graphs, and lower thresholds will tend to produce larger, more densely connected graphs.

### Section 2: Nonlinear effects of the threshold choice

Figure 2 shows the percentage of nodes in the largest component as a function of the choice of threshold, for each of the three treatments. To facilitate comparison with past work on this data set (Barberán *et al.* 2012), we include a dashed vertical line at a threshold of 0.36. This yields a network from the noise-added data of comparable size to this past work (*n=300*). The location of the noise transition in the green line (Δ), near a threshold of 0.30 represents a lower bound on reasonable choices of a threshold.

Across thresholds, the original data shows a relatively slow decline in the size of this largest component. Compared to the other treatments, which better eliminate spurious connections, this slow decline is clearly an artifact of the presence of many false positives in the network. By applying the Spearman correction or that correction and the null distribution from permutations, the largest component shrinks much more quickly. The difference between the treated lines and the original data illustrates the dramatic extent to which not controlling for these statistical artifacts can alter the extracted structure of the species interaction network. A further observation is that the smooth variation of the noise-added data treatment indicates that there is no obviously best choice for a threshold, except somewhere close to but slightly above the noise transition.

This finding illustrates the complexities that arise when using a threshold to extract a network from a correlation matrix, and suggests that a particular choice requires some justification or at least a robustness analysis to demonstrate that scientific conclusions do not depend sensitively on that choice. From a data analysis perspective, we would preserve the most ecological signal by not applying a threshold and instead using the correlation scores as weights for edges in a fully connected or complete graph (Thomas & Blitzstein 2011). However, many common network analysis techniques do not generalize to weighted complete networks, or such methods have not yet been developed. As a result, thresholding may be necessary to address certain classes of ecological questions.

To further illustrate the impact of threshold choice on the structure of the induced network, we measured five standard network summary statistics as a function of threshold choice. These summary statistics are (i) the average degree, (ii) the average path length, (iii) the diameter, which is the maximal-length shortest path among any pair of nodes, (iv) the modularity, which quantifies the extent to which nodes cluster into groups, with more edges occurring inside groups than expected at random, and (v) the clustering coefficient.

If the functional relationship between threshold and network statistic were constant or linear, the particular choice of threshold is less likely to impact scientific conclusions that depend on its particular value. For all five of these measures, however, we find a nonlinear relationship between the measure and the choice of threshold. That is, the structure of the network does not change smoothly, and different threshold choices can lead to very different patterns of connectivity within the network (Figure 3).

**Fig 3:**
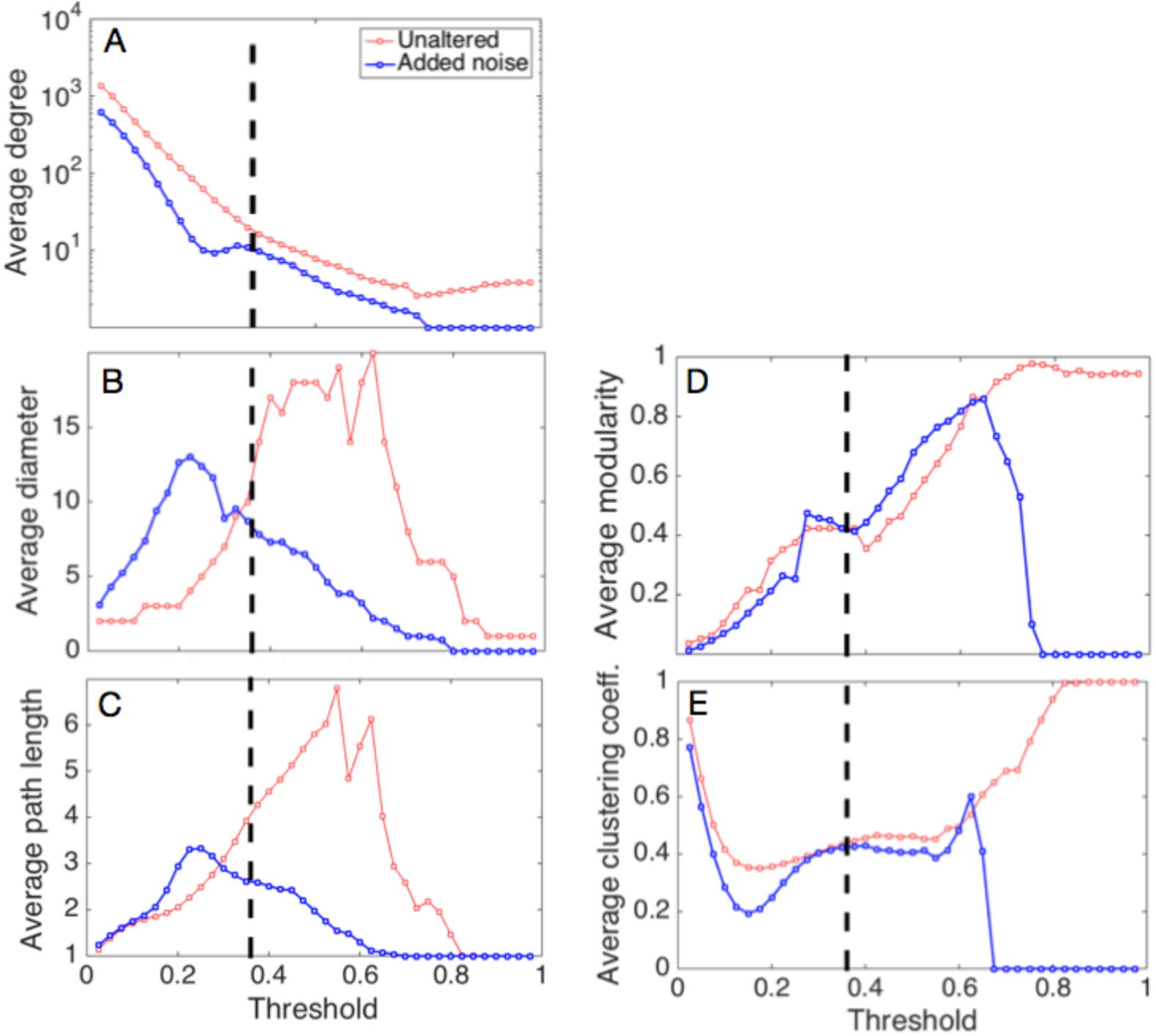
Network properties vary as a function of threshold. This figure shows the change of network properties as the similarity score threshold varies between 0 and 1. The red lines represent the unaltered abundance data; the blue lines represent the noise-added data to correct rank ties. The vertical line at 0.36 is the same threshold used in Figure 2. Panels correspond to the following properties: (A) average degree, (B) diameter, (C) average path length, (D) maximum modularity, and (E) clustering coefficient.

For instance, even the average degree of this network exhibits a surprisingly nonlinear pattern across thresholds (Figure 3A). The non-monotonicity, illustrated by the bump around a threshold of 0.35, results from the convention of discarding nodes with no connections. Thus, as the threshold increases, more of these nodes are created and then excluded, which allows the average degree to increase again as the giant component shrinks but the connectivity of its nodes stays relatively steady. (The average degree touches the x-axis at a threshold of 0.75; when singletons are included, this transition occurs around a threshold of 0.40 (Supp. Fig. 1).)

Similar patterns appear for the average and maximal path length (Figs. 3B and 3C). At lower thresholds, the network is relatively dense, making short paths among nodes plentiful. As the threshold increases, edges are removed, which makes the largest component sparser and increases path lengths. Finally, both measures decline above a threshold of 0.25 as the size of the largest component itself begins to shrink, which shortens path lengths again.

As the threshold increases, the largest component becomes sparser and the estimated maximum modularity score also increases (Figure 3D), implying the existence groups of nodes with relatively high internal connectivity (Clauset *et al.* 2004). This property deviates from a simple linear increase between threshold values of 0.25 and 0.38. At higher values of the threshold, the largest component breaks up into small but fully connected subgraphs, which have the highest possible marginal contributions to modularity. However, for very high threshold values, the average degree falls below 1 and the network is composed primarily of disconnected edges, which yields a modularity score of 0.

Because very low thresholds produce very dense networks, the clustering coefficient (Figure 3E) is initially very high, but decreases quickly. Interestingly, and unlike the other network statistics on this data set, the clustering coefficient stabilizes across intermediate choices of thresholds, even as other network statistics are still changing. As with the behavior of modularity, the clustering coefficient rises quickly and then falls to 0 as the network crosses from being composed primarily of disconnected triangles and edges to being composed entirely of disconnected edges.

The nonlinear dependence of the structure of the extracted network on the threshold applied to the correlation matrix demonstrates the importance of performing robustness analyses in this setting. Higher thresholds tend to naturally produce networks with many small components, high modularity and shorter path lengths. Lower thresholds tend to produce a large component, often with lower modularity scores. The threshold at which the transition between these two regimes occurs is likely to be data dependent, and thus should be quantified in order to clarify the confounding role that network size and density have on other network measures.

### Choosing a threshold for significant interactions

If there existed a labeled data set, such as fully-defined microbial communities where every individual microbial cell had fully sequenced 16S ribosomal RNA, we could train a machine learning model to choose the threshold that best balances false positive (spurious) links against false negative (missing) links, when those communities are sampled. However, it is typically impractical to fully characterize the taxa that make up an *in vivo* microbial community. Thus, in practice, choosing an intermediate value for the threshold is a reasonable strategy. The threshold should be large enough to be above the noise transition (Figure 2, green line), but small enough that the network is not mostly disconnected. However, because of the nonlinear relationships between network structure and threshold choice, a robustness analysis should always be performed in order to determine whether a particular conclusion depends sensitively on which intermediate threshold is chosen.

### Section 3: Measuring non-random network structure

Given a choice of threshold and the corresponding network derived from corrected Spearman correlation scores, we can now ask whether the distribution of the network’s links represents non-random patterns. We use a second class of null models to find statistically significant properties of the derived network by controlling for connectivity. The two models in this class will allow us to distinguish whether a particular pattern in the distribution of edges across the network is likely due to chance.

The first null model is the Erdős–Rényi random graph, which preserves the average degree of the derived network while removing any taxonomic information from the nodes (Erdős & Rényi 1960, Kara *et al.* 2013). This model is sometimes denoted *G(n,p)*, where *n* is the number of nodes and *p = <K>/(n-1)*, where the mean degree *<K> = 2m/n* is the probability that any pair of vertices is connected and where *m* is the number of edges in the derived network. Drawing a large number random graphs from this model (e.g., 2000 graphs) allows us to numerically estimate a null distribution for any network property, while controlling only for the average degree of a node.

The second null model in this class is a Chung-Lu random graph model (Chung & Lu 2002) where we prohibit self-loops (an edge *(i,i)* for some node *i*). Like the Erdős– Rényi model, a Chung-Lu model starts with the same number of nodes as the derived network. Rather than giving each edge equal probability, this model preserves the expected degree sequence by making the probability of an edge between two nodes proportional to the product of their expected degrees. Specifically, the probability of an edge between nodes *i* and *j* is *Pi,j = (ki * kj) / 2m−1,* where *ki* is the degree of node *i* in the derived network. This model is similar to the popular configuration model (Molloy & Reed 1995), but like the Erdős–Rényi model, it only produces simple networks, i.e., those without self-loops or multiple connections between the same pair of nodes. As before, drawing a large number of random graphs from this model allows us to numerically estimate a null distribution for the same network properties of interest, but now controlling for the average degree of a node and the degree distribution across nodes.

To illustrate how these models can be used to distinguish plausible structural patterns from those generated by chance, we apply them to the soil microbe network extracted in the previous section from the corrected Spearman scores. The derived network has about *n* = 268 nodes and *m* = 1730 edges; the precise numbers vary depending on the noise addition step. We then compare the null vs. the derived network’s distributions for (i) mean path length, (ii) modularity, (iii) diameter, and (iv) clustering coefficient. Both null models are parameterized to match the mean degree and thus the random graphs match the derived network on that measure by design.

Both path length and diameter are slightly elevated in the networks derived from the corrected Spearman data compared to the null models (Figures 4A-B). The average path length is 2.935 ± 0.052 for the corrected data, compared with 2.611 ± 0.019 for Erdős– Rényi and 2.604 ± 0.040 for Chung-Lu. Similarly, the average diameter for the corrected data is 7.411 ± 0.874, compared with 4.114 ± 0.318 in the Erdős–Rényi and 5.582 ± 0.544 for the Chung-Lu models. These differences are statistically significant, although the effect size is small. That is, the extracted microbial interaction networks are only less compact than we would expect if edges were distributed at random.

**Fig 4:**
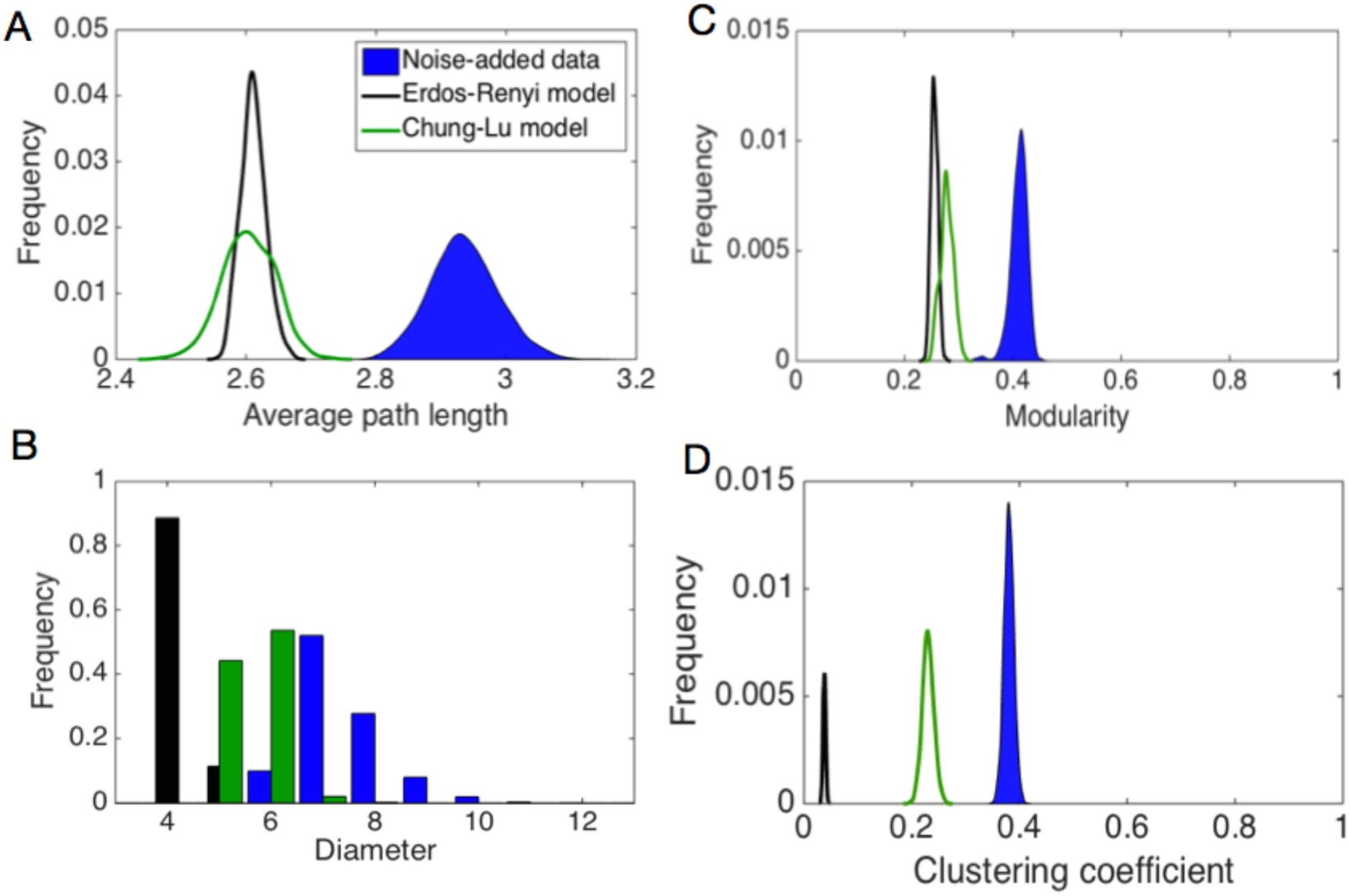
Network properties compared with null network models with fixed connectivity. Distributions of network properties across observed data and null models from the second class of models: Erdős–Rényi and Chung-Lu. The observed data is graphed as blue in each plot. Panels show the following properties: (A) average path length, (B) diameter, (C) modularity, and (D) clustering coefficient.

Similarly, the modularity scores (Figure 4C) are higher in the derived network compared to those of the null models. The modularity is 0.415 ± 0.014 for the derived network, while it is 0.217 ± 0.005 for the Erdős–Rényi model, and 0.280 ± 0.012 for the Chung-Lu model. For these null models, the observed modularity scores are highly statistically significant, and thus may represent a true ecological signal. However, as we observed in the previous section, the modularity score is highly dependent on the choice of threshold. For instance, under a threshold of 0.25 instead of 0.36, the difference in modularity scores between the Chung-Lu null model and the derived network vanishes (both are approximately 0.299). As such, the significance of the modularity score should be interpreted cautiously.

Compared to both null models, the derived network has a substantially higher clustering coefficient (Figure 4D), which is similar to the scores observed in social networks (Newman 2012; page 237). The clustering coefficient for the derived network is 0.380 ± 0.009, while it is 0.038 ± 0.002 for Erdős–Rényi random graphs and 0.230 ± 0.010 for Chung-Lu random graphs. The difference in null distributions indicates that about half of the value of the observed clustering coefficient can be explained as an artifact of heterogeneous degree structure, which the Chung-Lu model captures but the Erdős–Rényi model does not. This suggests that microbial communities are enriched in three-way interactions (triangles) and these represent potentially ecologically meaningful functional relationships among triplets of OTUs.

### The consensus network

Because the network properties of the derived network appear statistically significant relative to our random graph null models, we can now construct and interpret a “consensus network,” which contains every pairwise interaction that is present in at least 90% of the Monte Carlo samples. This consensus network is composed of 158 nodes and 787 edges. A simple but scientifically interesting question we may address with this network is whether microbes tend to co-occur with others in the same phylum. A positive signal of this assortative mixing pattern (Newman 2003) would suggest a phylogenetic structuring for niche preferences or potential synergistic relationships within phyla (Barberán *et al.* 2012).

However, we see little evidence for this hypothesis, finding instead that soil microbes are not more likely to co-occur with taxa within phyla rather than across phyla (Figure 5). Specifically, the number of edges between two phyla appears roughly proportional to the number of taxa in both phyla, exactly as we would expect if such co-occurrences were largely due to chance. As an additional check, we calculate the fraction of edges that connect each phylum (Table 1). This enables us to investigate the potential heterogeneous mixing of phyla. We observe that Acidobacteria and Proteobacteria have the highest proportions of within-phylum edges, so these phyla are most likely to co-occur with species within their respective phyla, when we enforce that clusters must correspond to phyla. But the modularity of this network, which provides a quantitative measure of assortativity among categorical labels on nodes (in this case, phyla), we find a score of 0.0745 – much lower than the estimated maximal modularity when nodes are allowed to mix independently of their phyla label (Fig. 4C). That is, non-phylogenetic factors dominate the structure of OTUs interactions in this data set.

**Fig 5:**
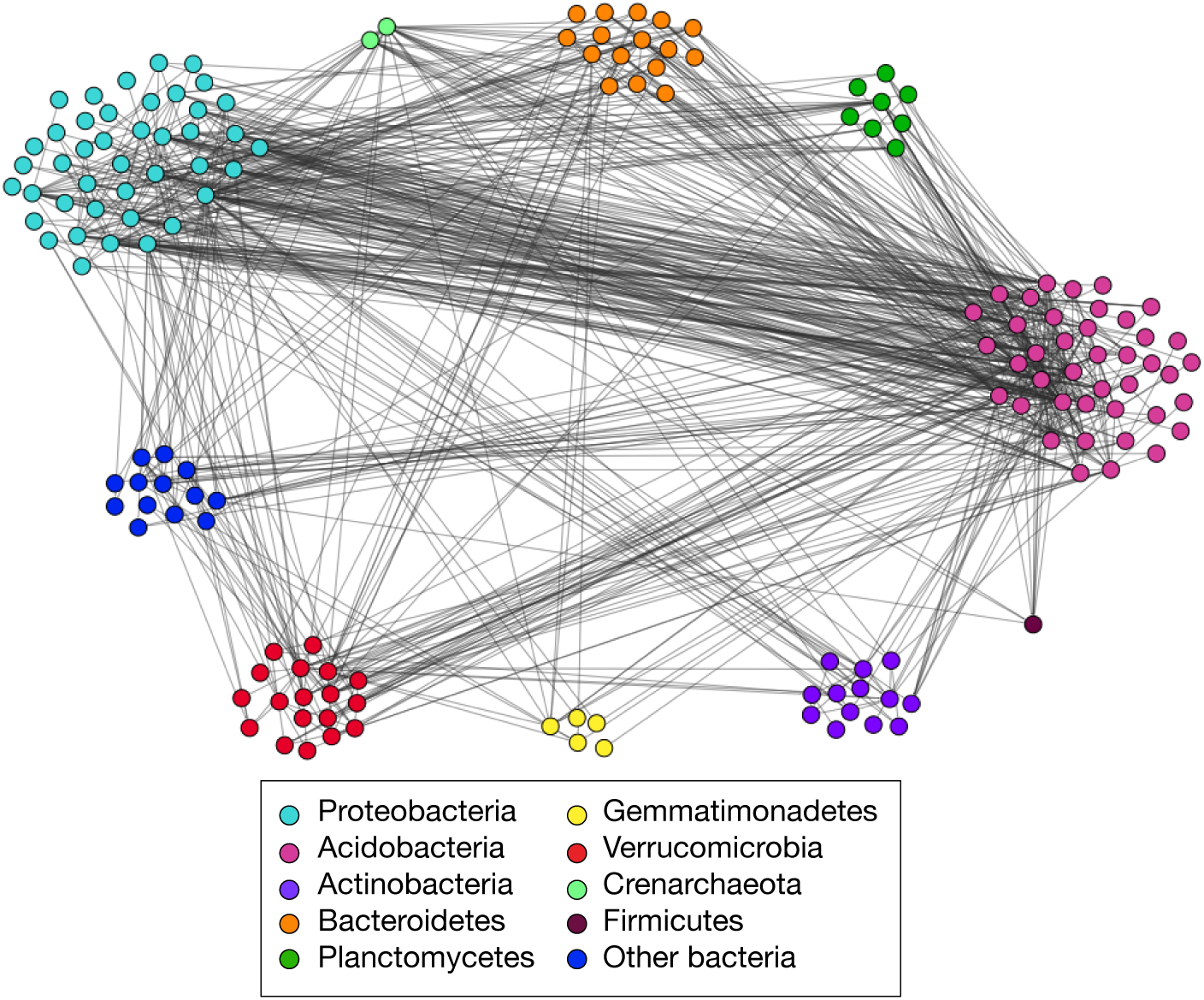
Consensus network of edges, organized by phylum. Edges in this figure are present in 90% of Monte Carlo simulations of noise addition.

**Table 1:** Fraction of edges connecting clusters based on phylum identity. Fraction of edges connecting pairs of OTUs across nine phyla (or “other”, for OTUs that don’t map to a known phylum). Edges connecting OTUs from the same phylum are highlighted. Note that this table is symmetric because edges are undirected.

|  | Acido | Actino | Other | Bacteroid | Crenarch | Firmicutes | Gemma | Plancto | Proteo | Verruco | All |
| --- | --- | --- | --- | --- | --- | --- | --- | --- | --- | --- | --- |
| Acido | **0.172** | 0.010 | 0.014 | 0.029 | 0.010 | 0.004 | 0.004 | 0.019 | 0.098 | 0.027 | 0.388 |
| Actino | 0.010 | **0.017** | 0.000 | 0.000 | 0.002 | 0.000 | 0.000 | 0.000 | 0.010 | 0.003 | 0.041 |
| Other | 0.014 | 0.000 | **0.004** | 0.002 | 0.002 | 0.001 | 0.001 | 0.001 | 0.011 | 0.008 | 0.043 |
| Bacteroid | 0.029 | 0.000 | 0.002 | **0.014** | 0.005 | 0.000 | 0.002 | 0.000 | 0.021 | 0.006 | 0.078 |
| Crenarch | 0.010 | 0.002 | 0.002 | 0.005 | **0.000** | 0.000 | 0.001 | 0.001 | 0.005 | 0.001 | 0.027 |
| Firmicutes | 0.004 | 0.000 | 0.001 | 0.000 | 0.000 | **0.000** | 0.000 | 0.000 | 0.001 | 0.000 | 0.006 |
| Gemma | 0.004 | 0.000 | 0.001 | 0.002 | 0.001 | 0.000 | **0.001** | 0.000 | 0.002 | 0.001 | 0.013 |
| Plancto | 0.019 | 0.000 | 0.001 | 0.000 | 0.001 | 0.000 | 0.000 | **0.000** | 0.014 | 0.001 | 0.036 |
| Proteo | 0.098 | 0.010 | 0.011 | 0.021 | 0.005 | 0.001 | 0.002 | 0.014 | **0.110** | 0.010 | 0.281 |
| Verruco | 0.027 | 0.003 | 0.008 | 0.006 | 0.001 | 0.000 | 0.001 | 0.001 | 0.010 | **0.031** | 0.087 |
| All | 0.388 | 0.041 | 0.043 | 0.078 | 0.027 | 0.006 | 0.013 | 0.036 | 0.281 | 0.087 |  |

## Discussion

A key step in better understanding the complex structure and function of microbial ecosystems is identifying the ecologically meaningful interactions among microbes. Distinguishing spurious interactions from real interactions is a key step in this process. However, common approaches in this setting can contaminate the extracted network with statistical artifacts that may confound ecological interpretation. Here, we have developed and demonstrated simple and appropriate null models for addressing this question at both the network extraction and the analysis steps, and we used them to reanalyze a previously studied large soil microbiome data set.

After adding noise to the sparse OTU abundance data, we examined in detail the difficulty of choosing a similarity threshold. Since network analysis depends on this initial network derivation step, a conservative approach would test whether conclusions about the network hold (or the same pattern appears) across a range of reasonable threshold choices. In practice, we suggest choosing a threshold slightly above the noise transition produced by the permutation test, and well below the point where the network breaks up into small, disconnected components. An interesting line of future work would examine the efficacy of supervised learning techniques from machine learning to automatically choose a threshold that optimizes some downstream performance measure (De Choudhury *et al.* 2010), e.g., likelihood of the extracted network under a probabilistic generative model like the stochastic block model (Karrer & Newman 2011).

Next, we used null models that preserve network connectivity to investigate the variation in network measures. We did find slight but statistically significant elevation of average path lengths and diameters in the derived network. One interpretation is that microbial communities in soil are robust to environmental perturbations and have evolved to recover or maintain structural stability amidst disturbances. Combining future research on different microbial communities, such as the human gut microbiome, with this type of network analysis would help clarify the role of average path length and diameter (if any) in community robustness, e.g. after the administration of antibiotics.

We discovered that the clustering coefficient was higher in the derived network compared to the network null models and that the score remained consistent across a range of intermediate threshold values. The elevated clustering coefficient may imply that habitat filtering is playing an important role in the distribution and abundance of OTUs in the soil. However, more research is needed to incorporate metabolic data or other functional predictors into the model. Levy and Borenstein (2012) have shown that in the human microbiome, co-occurrence is more often found in metabolically competitive species than in metabolically complementary species -- evidence that community assembly is best explained by habitat filtering in the human gut. Similarly, Goberna *et al.* (2014) also found that phylogenetic clustering was stronger in habitats where competitive traits prevailed (i.e., in areas with high resource availability). Future analysis of soil microbes should focus on metabolic competition and complementarity, especially within OTU triads, to determine whether the elevated triad occurrence corresponds to a specific community assembly mechanism (e.g., Pholchan *et al.* 2013, Coyte *et al.* 2015). Future inquiry should focus on whether elevated clustering coefficients are also present in networks derived from freshwater, marine, and human microbiome samples.

We also discovered elevated maximum modularity scores in the derived network compared to the null models. Higher modularity has been interpreted as corresponding to greater niche partitioning (Faust & Raes 2012, Montoya *et al.* 2015). Further analysis ofmetabolicfunctionsofOTUsshouldinvestigatewhetherthehighest-scoring modularity partitions indicate true functional niches, wherein OTUs are more likely to co-occur with OTUs in their own group than with OTUs in outside groups. For example, gene expression data can be compared within and across the proposed functional niches to identify shared or related metabolic functions (Levy & Borenstein 2013). Future work may glean more from co-occurrence networks that focus on the level of genes, rather than OTUs, which will become increasingly informative as more microbial genomes are fully sequenced.

The consensus network was composed of 50% generalist OTUs and 50% OTUs that were neither generalists nor specialists. The 79 generalist OTUs were identified based on appearing in more than 80 locations. The other half of the OTUs were neither generalists, nor specialists which appear in fewer than 10 sites with more than 18 sequences on average (Barberán *et al.* 2012). While no specialists appeared in the consensus network, only 17 OTUs out of the 1577 total OTUS were identified as specialists; given that about 10% of OTUs appeared in the consensus network, the expected number of specialists in the consensus network would be 1.7. It is not possible from this study to distinguish whether consensus networks are inherently biased against specialists or whether there was simply not enough data in this sample to distinguish specialists from noise. We do observe, however, that 76% of generalists (79 out of 104) are included in the consensus network.

The consensus network’s strong modularity score may be due to the relative concentration of generalists (Barberán *et al.* 2014). This might also explain why the optimal partitioning of the consensus network did not correspond with phylogeny, which was unexpected. The consensus network partitioning contrasts with basic assumptions that ecological functions and niches are phylogenetically conserved (Philippot *et al.* 2010). However, other recent work (Langille et al. 2013, Martiny et al. 2013) shows that while complex traits and housekeeping genes are generally deeply conserved, other functional traits like assimilation of carbon sources are broadly dispersed with respect to phylogeny. More work is required to identify the degree and manner in which functional diversity structures real co-occurrence in the soil microbiome.

The consensus network incorporates taxonomic information into the microbial interaction networks, allowing us to use 16S rRNA sequence similarity to evaluate the network structure. How best to incorporate that phylogenetic information is another area of active research (Agler *et al.* 2016). Previous research has shown that using lower binning thresholds for OTU identification does not reveal more about microbial interactions, suggesting that even relatively broad binning strategies can be useful for gaining ecological insight (Knights *et al.* 2011, Faust & Raes 2012). However, other authors recommend using the highest possible similarity threshold (Berry & Widder 2014). Future research should continue to address the phylogenetic information we have about OTUs and how that data can be incorporated into identifying real ecologial interactions (O’Dwyer *et al.* 2012).

Many recent microbial association studies have focused on problems with analyzing compositional data. For instance, several studies point out that compositional effects are a concern when there are big differences in component sizes (Yang *et al.* 2016) or when there are relatively few components (Ban *et al.* 2015). These problems are more prevalent in marine metagenomics samples or host-associated microbes, but less for the soil microbiome. We argue that using a relatively simple and nonparametric similarity measure such as Spearman correlation coefficient can prevent the imposition of preexisting notions about how taxa are distributed and how they interact. Compared to other techniques, Spearman correlation coefficients are also efficient to calculate, a problem acknowledged in the mLDM algorithm by its authors (Yang *et al.* 2016). For data not derived from the soil microbiome, the suggested approaches for compositional data could be used in conjunction with the network derivation methods described here.

In general, analyses of OTU-location matrices have uncertain scientific value as long as we lack large sets of empirically validated OTU-OTU interactions by which to evaluate network extraction methods. One possible remedy for this would be to remove some fraction of observed edges from the inferred network and use predictive modeling to identify the most probable missing edges. This type of link prediction has been used in other contexts where the observation of the network is incomplete or error-prone (Goldberg & Roth 2003), or where the network is changing, as in evolving social networks (Liben-Nowell & Kleinberg 2007). A generative link-prediction model allows us to test the degree to which our assumptions about the underlying structure of the system are correct (Clauset *et al.* 2008, Guimerá & Sales-Pardo 2009).

Future research should apply different models to recover community structure. The bipartite stochastic block model (Larremore *et al.* 2014) offers a compelling alternative to clustering OTUs based on their similarities across locations. That is, instead of converting the abundance matrix into a similarity matrix and applying an arbitrary threshold, this model operates directly on the OTU-location matrix, obtaining both a clustering of OTUs, a clustering of locations, and a mixing matrix that describes how OTU groups interact with location groups. By operating on the original OTU-location data, this approach would reduce the number and strength of assumptions used in the analysis of such data. This model would be useful for finding OTUs that co-occur and thus may be ecologically interacting, though it is defined only for occurrence data rather than abundance data. To include sequence abundances, the weighted stochastic block model (Aicher *et al.* 2015) could be used to directly analyze the OTU abundance values, without having to choose a threshold. For the task of clustering OTUs, these community detection methods are a promising set of tools.

While the approach we have outlined for testing different network derivation thresholds and evaluating null model connectivity has been applied to microbial abundance data, it can also be applied across other biological network analyses. Sparse data sets appear in a wide variety of biological settings, from eukaryotic environmental DNA surveys (e.g., Stoeck *et al.* 2010) and gene co-expression networks. The issues of threshold choice and appropriate null model selection are relevant across all disciplines which use network science. Utilizing the appropriate statistical approaches will allow researchers to draw stronger conclusions about correlation data, while leveraging the quantitative tools from network science accurately.

## Materials and Methods

Figure 6 illustrates the data analysis procedure used in this research.

**Fig 6:**
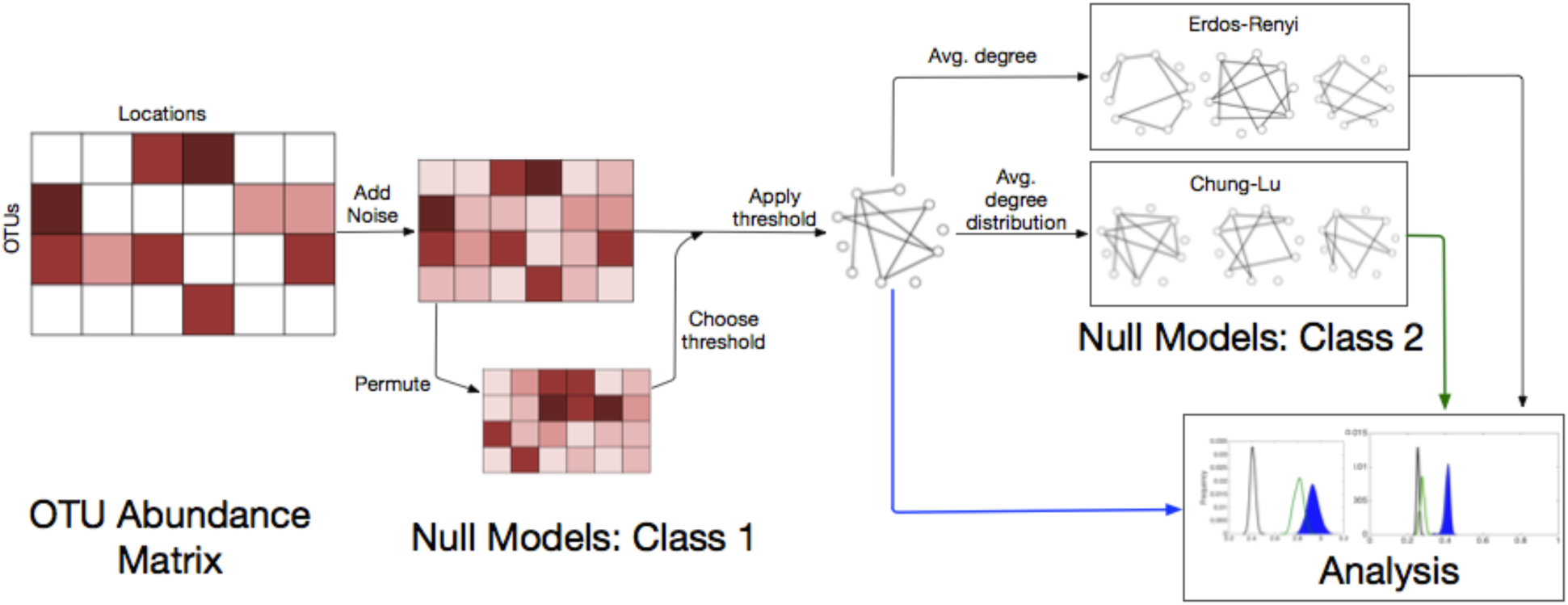
Data analysis procedure for OTU abundance matrices. We start with the OTU abundance matrix of *N* OTUs at *L* different locations. In the first class of null models, noise is added to every entry of the matrix. Additionally, the noise-added matrix is permuted; the distribution of similarity scores in the permuted matrix is used to set the lower bound for the threshold. Next, the threshold is applied to derive the observed network. This network is used to construct the second class of null models, Erdős– Rényi, based on the average degree, and the Chung-Lu model, based on the average degree distribution. Finally, the null network properties are compared to the observed network properties in the analysis step.

### Soil data

The data set used in this experiment was acquired from previous work. Lauber *et al.* (2009) acquired the bacteria and archaea data by pyrosequencing soil samples from locations across North and South America. Their data set covers 151 sampling sites and 4088 unique OTUs, binned at 90% similarity (for explanation of the choice of 90% similarity for binning, see Barberán *et al.* 2012). The data excludes OTUs with fewer than 5 sequences across all sampling sites, decreasing the number of OTUs to 1577. We use Spearman rank correlation coefficients to evaluate similarities between pairs of OTUs based on their abundance patterns. For each OTU, Spearman converts a vector of abundances into a vector of ranks, from largest to smallest. When there are identical abundance values in several locations for a given OTU, the corresponding locations in the rank vector are assigned the average rank for all tied entries. Given a pair of such rank vectors x and y, the Spearman rank correlation coefficient is given by:

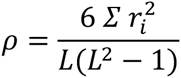

where *ri*=*xi* - *yi* is the difference between ranks between OTU *x* and OTU *y* in location *i*, and where *L* is the number of locations.

### Random noise addition

Rather than allowing for ties among Spearman ranks, we correct for sparsity in the OTU abundance matrix ***A*** by adding noise to every OTU × location entry. We draw N × L entries from a uniform distribution, *U*([0,1]), creating an N × L matrix *rand(N,L).* To ensure that we are breaking ties without reversing any true orderings, we adjust the random values to be several orders of magnitude smaller than the minimum difference between entries in ***A***:

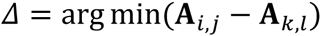

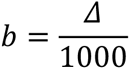

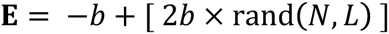

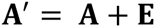

To ensure that the configurations of equally likely location ranks were well sampled, we repeated the noise addition steps 2000 times to generate a distribution of plausible interaction networks.

### Random matrix permutations

The most basic null model is the element-wise permutation of the OTU abundance matrix. We chose a uniformly random permutation of the entries in the OTU abundance matrix while maintaining the background distribution of abundances from which the values were sampled. The permuted data quickly transitions to having <1% of the OTUs in the largest component; this is where we set the lower bound for the similarity score threshold.

### Thresholding

The threshold that we use, 0.36, produces a network of approximately 300 nodes from the sparsity-corrected Spearman score data. This threshold was chosen to improve comparability between our results and those of past studies on the same data (Barberán *et al.* 2012). It is also similar to the threshold used by Friedman and Alm (2012) of 0.30. We did not identify any additional quantitative guidelines for threshold choice in other studies.

### Network derivation

The network was derived by defining edges as connections between pairs of OTUs with a *ρ* value greater than the absolute value of the chosen threshold. Nodes with no edges (also known as singletons) were omitted from the network, which is conventional in defining the network. Average degree, average path length, and diameter were calculated following the definitions in Newman (2010). Average degree is given by:

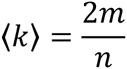

where *m* is the number of edges and *n* is the number of nodes. The average path length is given by:

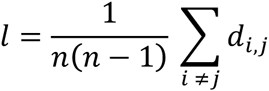

where *d_i,j_* is the shortest path between nodes *i* and *j* (this is different from the *d_i_* values used for the Spearman rank calculation). Diameter is the maximum value of *d_i,j_* across all pairs of nodes in the network (i.e., the longest shortest path).

The clustering coefficient is defined as the global proportion of open triangles that are closed by a third edge. We find all open triangles (i.e., paths of length 2) by taking the dot product of the derived network’s adjacency matrix ***Q*** with itself. Since this is an undirected graph, we analyze the upper triangle of the matrix only, not including the diagonal. Next, to find the proportion of length-two paths traversing three nodes that are also closed triangles, we multiply the upper triangle by the original matrix ***Q***, element-wise. The clustering coefficient *c* is the fraction of open triangles that contain a third edge to close the triad:

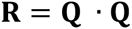

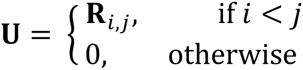

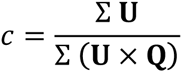

The maximum modularity was calculated using the using the popular greedy agglomerative algorithm of Clauset, Newman and Moore (2004). This algorithm begins with all nodes in their own group and then repeatedly merges the pair of groups that maximizes the marginal improvement in the modularity score until only one group remains. It then reports the maximum modularity value Q and the corresponding grouping of nodes D that it traversed in this sequence. We used the implementation of the algorithm in the igraph package in R (Csardi & Nepusz 2006).

### Class 2: Null network models

Erdős–Rényi random graphs were created based on the average degree of the derived network. Given an average degree of 11.64 and 300 nodes:

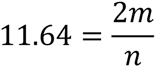

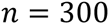

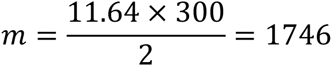

Once we had calculated the number of edges that we wanted in order to produce similar average degrees to the real data, we used the 300 × 300 adjacency matrix **T** to determine the correct threshold *p*:

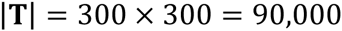

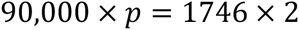

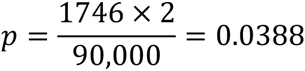

To derive the Erdős–Rényi random graphs, we generated a 300 × 300 uniform random matrix. We thresholded the upper triangle of the matrix at 1 − 0.0388 = 0.9612 and reflected it across the diagonal. All entries on the diagonal were set to 0. This approach is consistent with the mathematical definition of Erdős–Rényi random graphs, where edges are randomly chosen for all pairs. Thus, the Erdős–Rényi graph has no self-loops or multi-edges, as each pair is handled once.

For the second null model on the derived network, we used a modified Chung-Lu model to produce edges between nodes while preserving the expected degree distribution. The probability that an edge exists between OTU *i* and OTU *j* is given by,

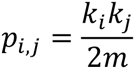

where *ki* is the degree of node *i* in the derived network. To generate a single Chung-Lu network we set *n* equal to the number of nodes in the observed network. Then, for each pair of nodes (*i, j*), we picked a uniform random number between 0 and 1. If the random number was between 0 and *pi,j* -- which is proportional to the product of their degrees –we created an edge connecting nodes *i* and *j* in the Chung-Lu network. We repeated this method 2000 times to generate a distribution of Chung-Lu random graphs.

### Consensus network

To derive the consensus network, we applied 2000 Monte Carlo runs to the corrected Spearman data at the 0.36 threshold. We included edges that appeared in 90% of the trials to produce the consensus network. We visualized networks using the software Gephi (Bastian *et al.* 2009). Nodes were color-coded by phylum.

### Code

All code for processing the data, applying null models, deriving networks, and measuring network properties are publically available on GitHub. The repository is saved under nkinboulder/MicrobeCommunities. The code for this project was written in Matlab and it is extensively commented for clarity.

## Acknowledgments

The authors thank Noah Fierer, Diana Nemergut, Christopher Aicher, Lauren Shoemaker, and Andrea Berardi for helpful conversations, and thank Noah Fierer for sharing data.

